# Hepatic gene replacement restores energy metabolism and doubles the survival in mouse model of GRACILE syndrome, a neonatal mitochondrial disease

**DOI:** 10.1101/2025.09.23.677965

**Authors:** Rishi Banerjee, Janne Purhonen, Nasrin Sultana, Christa Kietz, Vineta Fellman, Jukka Kallijärvi

## Abstract

Preclinical gene therapy studies of mitochondrial diseases remain limited due to their multisystemic manifestations and the scarcity of physiologically relevant animal models. Mutations in *BCS1L*, a nuclear gene encoding an assembly factor for mitochondrial complex III (CIII), are the most common cause of pediatric CIII deficiency. The most severe phenotype, GRACILE syndrome, is caused by a homozygous Finnish founder mutation (*c.A232G*, *p.S78G*). The corresponding *Bcs1l^p.S78G^* knock-in mouse model effectively recapitulates the human disease, with juvenile-onset hepatopathy, tubulopathy, growth restriction, segmental progeria, and short survival. Here, we performed recombinant adeno-associated virus (rAAV)-mediated gene replacement in this model, which features a postnatal presymptomatic window until weaning. A single intraperitoneal injection of rAAV encoding wild-type *Bcs1l* restored CIII assembly and activity in the liver, preventing hepatopathy. Hepatocyte-specific correction was sufficient to ameliorate hypoglycemia, improve growth, normalize systemic metabolism, and extend survival by nearly two-fold, despite persistent CIII deficiency in other tissues. Remarkably, liver-directed rescue also prevented skeletal muscle transcriptomic changes, particularly those linked to altered energy substrate utilization. These results underscore the central role of the liver in systemic energy homeostasis and growth regulation in multiorgan mitochondrial diseases and demonstrate the therapeutic potential of hepatocyte-directed gene replacement in phenotypes with prominent hepatopathy.

Graphical abstract

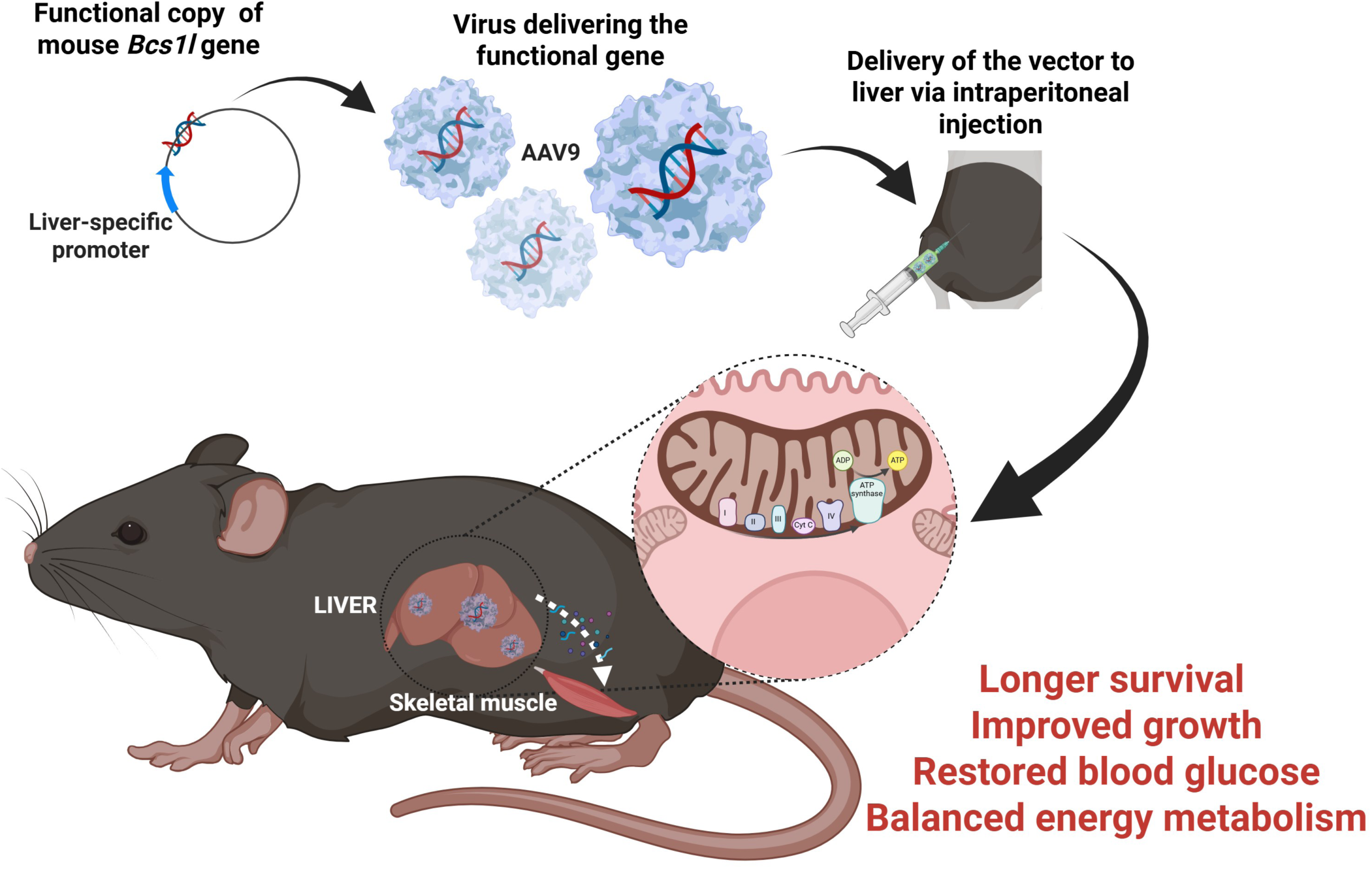

## Introduction

Mitochondrial disorders are a heterogeneous group of inherited metabolic conditions, often affecting oxidative phosphorylation (OXPHOS)^1,2^. They can arise from mutations in either mitochondrial or nuclear genes, manifest at any age, and involve almost any organ. However, most mitochondrial diseases are myopathies, encephalopathies, or encephalomyopathies. CIII deficiencies are relatively rare among them^3^. The most severe phenotype, GRACILE syndrome (**g**rowth **r**estriction, **a**minoaciduria, **c**holestasis, liver **i**ron overload, **l**actic acidosis, and **e**arly death), is caused by a homozygous *BCS1L* missense mutation (c.A232G, p.S78G)^4,5^. A knock-in mouse model carrying the GRACILE syndrome mutation recapitulates many features of the syndrome, including hepatopathy, kidney tubulopathy, and systemic metabolic crisis^6,7^. The liver is one of the most prominently affected organs in GRACILE syndrome patients and in the *Bcs1l^p.S78G^*mice. How the systemic phenotypes, such as growth restriction, loss of white adipose tissue, hypoglycemia, relate to the liver disease remains unclear^8^. In general, interorgan communication between the affected and unaffected organs in mitochondrial diseases is poorly understood, largely due to the lack of suitable physiologically relevant models^9,10^.

In gene therapy, the wild-type version of a mutated gene, or other therapeutic gene, is introduced into the affected tissues using suitable delivery strategies, typically to treat genetic conditions^11,12^. Among the available viral vectors for gene therapy, recombinant adeno-associated viruses (rAAVs) are widely used due to their safety and broad tissue tropism^13,14^. While efficient systemic gene delivery remains challenging, current technologies can target specific cells or tissues to achieve therapeutic effects. In clinical trials of genetic diseases, rAAVs have shown efficient gene delivery across a variety of organs and tissues, including the liver (ClinicalTrials.gov ID: NCT00377416, NCT02082860, NCT02484092), eye (ClinicalTrials.gov ID: NCT02946879, NCT03001310, NCT02781480) and brain (ClinicalTrials.gov ID: NCT00229736, NCT05603312, NCT00195143). Organ-targeted gene therapy has shown robust effects in some preclinical models of mitochondrial disorders^15–23^. Leber’s hereditary optic neuropathy (LHON) is, however, currently the only primary mitochondrial disease undergoing clinical trials for gene replacement therapy^24^. Liver-targeted gene therapy has rescued lethality in a zebrafish model of Leigh syndrome (mutated *LRPPRC*) and in a mouse model of mitochondrial neurogastrointestinal encephalomyopathy (mutated *TYMP)*^19,25,26^.

Here, we delivered rAAVs encoding wild-type mouse *Bcs1l* under a broadly active or hepatocyte-specific promoter^27^ to pre-symptomatic *Bcs1l^p.S78G^*mice. We used mice of the genotype *Bcs1l^p.S78^*;*mt-Cyb^p.D254N^* throughout this study because in this *Bcs1l^p.S78G^* mutant strain carrying a spontaneous *mt-Cyb^p.D254N^*variant^28^, the homozygotes become terminally ill at approximately 1 month of age, more closely resembling the neonatally lethal human disease and also allowing fast assessment of survival. As primary outcomes, we assessed growth and survival. We show that the rAAV-based gene replacement is safe and effective and that hepatocyte-specific rescue of CIII function is sufficient to ameliorate both hepatic and systemic manifestations and double the survival of the mice.

## Results

### Liver-specific rAAV-based gene therapy doubles the survival of CIII-deficient mice

The protocol we used for the rAAV administration and assessment of disease progression is shown in Fig. 1A. Hepatocyte-specific ApoE enhancer and α1-antitrypsin (AAT) promoter^29^ drove the expression of enhanced green fluorescent protein (EGFP), as a control, or mouse BCS1L, as a treatment. We refer to rAAVs carrying the *Bcs1l* or *EGFP* transgene as AAT-*Bcs1l* and AAT-*EGFP,* respectively. Fluorescence microscopy (Supplementary Fig. 1A) demonstrated effective transduction of the liver in the AAT-*EGFP*-injected mice, similar to previously published^30^. The mutant livers that received AAT-*Bcs1l* appeared visually as healthy as the WT livers one week after the injection (Fig. 1B). qPCR analysis of total and virally expressed *Bcs1l* at postnatal day 28 (P28) liver confirmed the viral expression of *Bcs1l*, which resulted in an approximately 15-fold increase in total *Bcs1l* mRNA in the liver (Fig. 1C).

**Figure 1.**
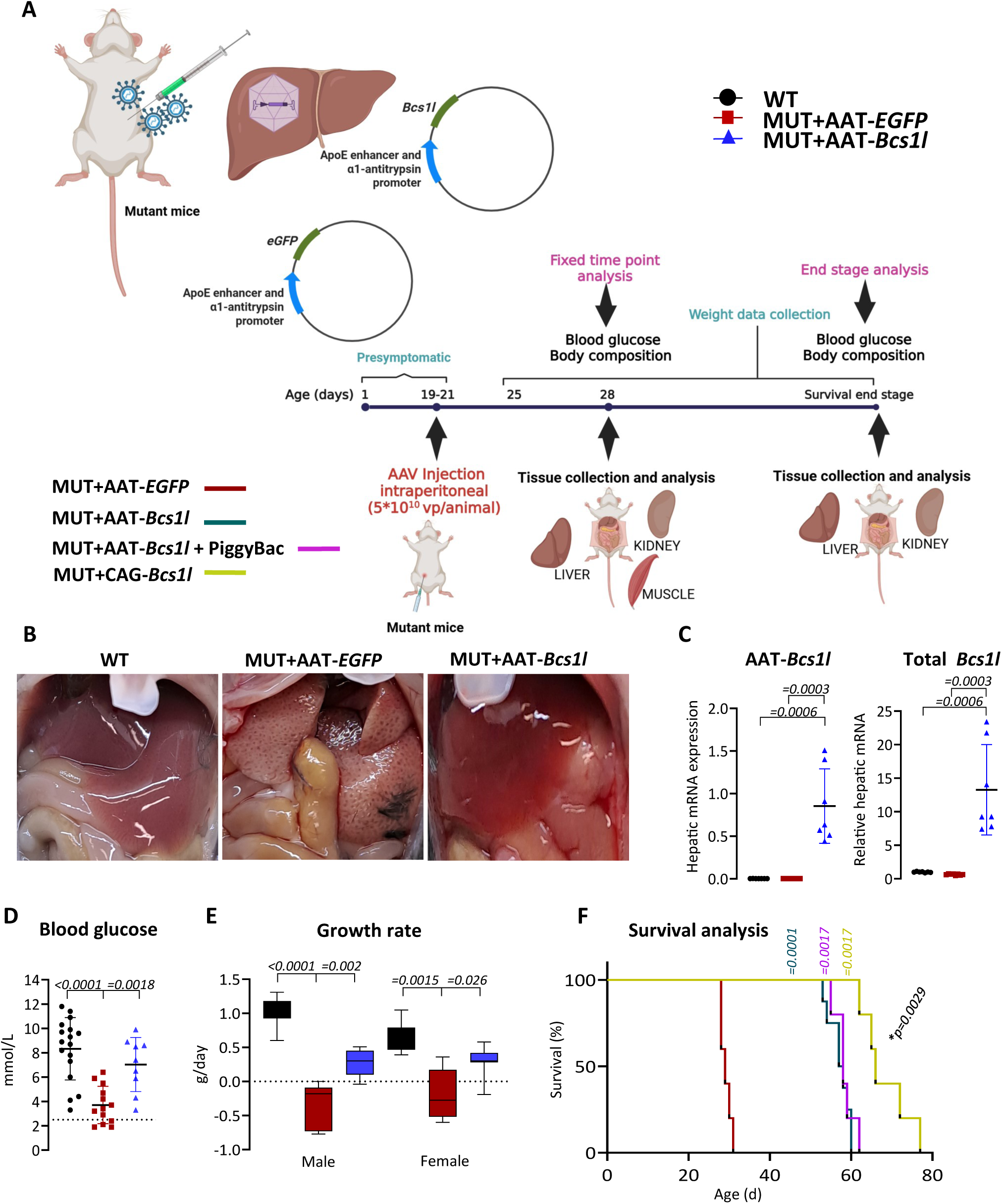
rAAV-based gene replacement rescues the growth and doubles the survival of CIII-deficient mice. A) Schematic presentation of the experimental setup and the timeline of the investigations. B) Appearance of the livers at P28. C) Virally expressed and total *Bcs1l* mRNA at P28 liver (*n*=7-8/group). D) Blood glucose levels at P28 (*n*=9-17/group). The dotted line indicates the critical level of glucose (<2.5mmol/L) predicting spontaneous death. E) Sex segregated growth rate of the mice from P25 to P28 (n=6-13/group). F) Survival curves of rAAV-*EGFP*-injected mutant mice and those of mutant mice injected with three different *Bcs1l*-expressing rAAVs (*n*=5-8/group). Statistics: Mann-Whitney U test (C). One-way ANOVA followed by the selected pairwise comparisons with Welch’s t-statistics (D and E) and log-rank test (Mantel-Cox) (F). *\*p*, comparison between survival of AAT*-Bcs1l* and CAG*-Bcs1l* group. The error bars stand for standard deviation. All data points derive from independent mice.

The *Bcs1l^p.S78G^*;*mt-Cyb^p.D254N^* mice succumb to metabolic crisis with extreme hypoglycemia at approximately one month of age (P30-35)^6,8,28^. At P28, AAT-*Bcs1l* significantly improved the low blood glucose, and, importantly, none of the treated mice showed blood glucose less than <2.5mmol/L (Fig. 1D). While AAT-*EGFP*-treated mutants lost weight after P25, the mutants treated with AAT-*Bcs1l* were able to grow (Fig. 1E). The AAT-*EGFP*-injected mutant mice reached the criteria of euthanasia between P28 and P30 (Fig. 1F). In contrast, the mutants injected with hepatocyte-specific rAAV-*Bcs1l* showed no signs of terminal deterioration or spontaneous deaths before the age of P53. Their median survival (criteria fulfilled for euthanasia) was 58 days (Fig. 1F). Towards the end of the extended survival, the glucose levels became critically low again (Supplementary Fig. 1B).

Being an episomal vector system, expression from rAAVs can dilute or fade away over time. Compared to the livers from P28 mice (Fig. 1C), *Bcs1l* mRNA expression indeed decreased by the end stage (Supplementary Fig. 1C). To assess if vector dilution accounted for the eventual deterioration of the AAT-*Bcs1l*-treated mice, we co-injected a PiggyBac transposase-encoding rAAVs with the AAT-*Bcs1l* virus for persistent long-term expression via genomic integration^29^. This strategy resulted in significantly higher end-stage hepatic *Bcs1l* expression (Supplementary Fig. 1C), but the survival did not extend further (Fig. 1F). This result suggests that the eventual deterioration was due to the disease progressing in the other affected organs. For further analyses of the survival panel, the AAT-*Bcs1l+*Piggybac transposase-injected mutants were used to ensure the hepatic expression of *Bcs1l* was not limiting. To interrogate the effects of extrahepatic transduction, we used a construct with the broadly active CAG promoter (CAG-*Bcs1l*) with an identical dose and intraperitoneal injection. The broader *Bcs1l* expression did increase the survival further, albeit relatively modestly (15%), to a median of 66 days (Fig. 1F). This vector also led to high-end-stage hepatic *Bcs1l* expression even without genomic integration (Supplementary Fig. 1C).

### Efficient restoration of hepatic CIII assembly and activity

Because we have previously extensively characterized the respiratory complex assembly and activities^6,28,31,32^ in the *Bcs1l^p.S78G^* and *Bcs1l^p.S78G^*;*mt-Cyb^p.D254N^* mice^28,32–35^, we only assessed CIII assembly and activity in the liver here. In isolated liver mitochondria, following AAT-*EGFP* treatment, blue native gel electrophoresis (BNGE) and immunoblot analyses showed a decreased amount of fully assembled CIII dimer (CIII_2_) based on the decreased UQCRFS1 (RISP) protein. Almost all of the residual fully assembled CIII_2_ was in the CI-CIII_2_ supercomplexes (SCs), with free CIII_2_ being absent. AAT-*Bcs1l* efficiently increased the levels of fully assembled CIII_2_ (Fig. 2A). In isolated mutant liver mitochondria, the mean CIII activity was 26% of WT. AAT-*Bcs1l* increased the mean CIII activity to 74% of WT (Fig. 2B) and improved hepatic ATP levels (Fig. 2C). We have previously estimated that symptoms appear at approximately 50% residual activity^28^. Even though the correction of CIII assembly and activity was partial, it still clearly exceeded the ∼50% threshold required to prevent disease onset.

**Figure 2.**
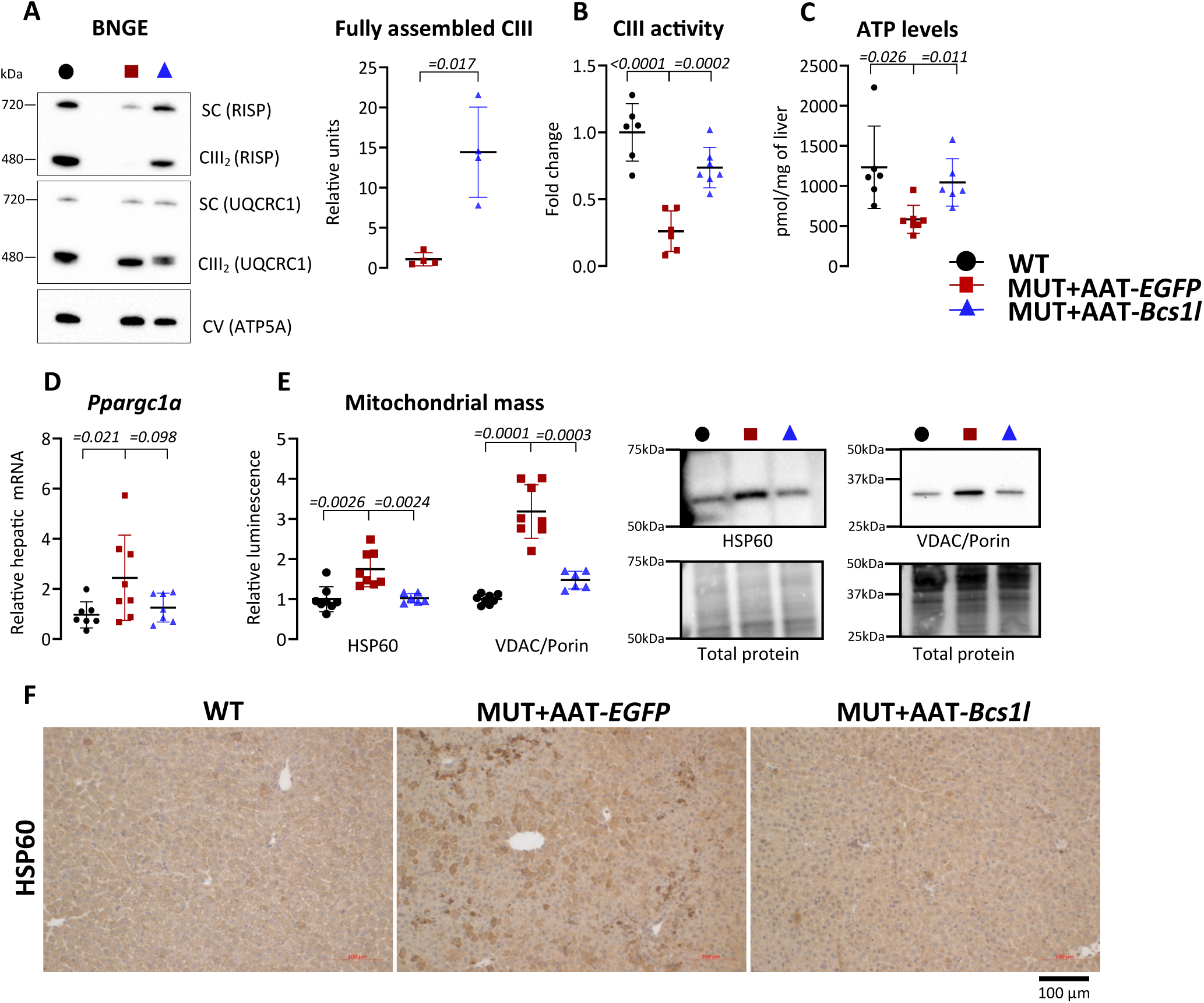
Hepatocyte-targeted gene replacement corrects hepatic CIII assembly and activity. A) Representative blue-native PAGE blot of UQCRFS1 (RISP) and UQCRC1 in free CIII_2_ and super complexes (SCs) from isolated P28 liver mitochondria, and RISP per total UQCRC1 ratio (*n*=4/group). B) CIII activity normalized by total protein in liver isolated mitochondria at P28 (*n* = 4/group). C) ATP in liver at P28 (*n* = 6-7/group). D) mRNA expression of *Ppargc1a (PGC-1α)*, from P28 liver (*n*=7-8/group). E) Western blot quantification and representative blots of HSP60 and VDAC1 as markers of mitochondrial mass from P28 liver lysates (*n*=6-8/group). F) Representative images of liver sections immunostained for the mitochondrial marker, HSP60. Statistics: Welch’s *t*-test (A), one-way ANOVA followed by the selected pairwise comparisons with Welch’s t-statistics (B-E). The error bars stand for standard deviation. All data points derive from independent mice.

The CIII deficiency leads to increased mitochondrial biogenesis and mass in the *Bcs1l^p.S78G^* livers^31,35^. Here, the mitochondrial biogenesis-driving key transcriptional regulator *Ppargc1a* (*Pgc-1α*) (Fig. 2D) and the mitochondrial mass markers VDAC1 and HSP60 were significantly increased in the untreated mutant livers but not in the AAT-*Bcs1l*-treated livers (Fig. 2E). Immunostaining of liver tissue sections showed frequent hepatocytes with irregular, abnormally strong HSP60 staining. The AAT-*Bcs1l* fully prevented the abnormal HSP60 staining pattern (Fig. 2F).

### Essentially complete prevention of liver pathology and correction of energy metabolism

Histopathological analysis of the P28 mutant mice showed hepatopathy characterized by expansion of portal areas, increased ductular reactions, and cell death (Fig. 3A-D). The AAT-*Bcs1l* fully prevented these changes, as well as the upregulation of the mitochondrial dysfunction-associated mitokine growth-differentiation factor 15 (*Gdf15*) (Fig. 3E). In the end stage of the survival, the hepatic ductular reactions had increased minutely (Supplementary Fig. 2A-D) and *Gdf15* expression was increased (Supplementary Fig. 2E), suggesting eventual incipient liver pathology despite the remaining WT *Bcs1l* expression.

**Figure 3.**
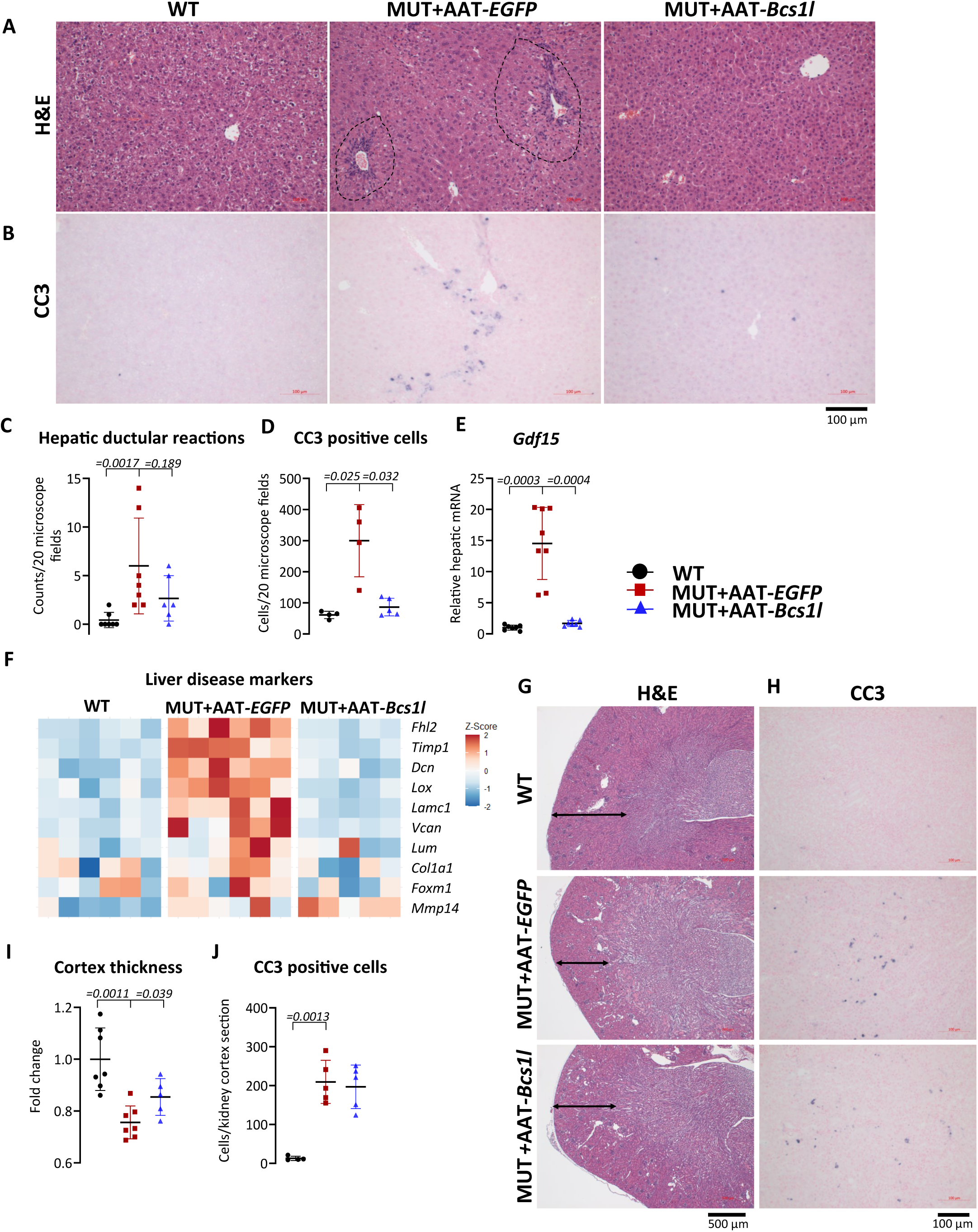
Gene therapy prevents mitochondrial hepatopathy and improves kidney cortex thickness in CIII-deficient mice. A and B) Representative images of H&E-stained liver sections (*n*=6-7/group), showing tissue morphology and expansion of portal areas (indicated by dotted lines), and liver sections immunostained for apoptotic cell marker, cleaved caspase 3 (CC3) (*n*=5/group) at P28. C and D) Quantification of hepatic ductular reactions and apoptotic cells from H&E-stained and CC3-immunostained sections, respectively. E) *Gdf15* mRNA expression from P28 liver (*n*=7-8/group). F) Heat map visualization of gene expression linked to liver disease (n=5-6/group). G and H) Representative images of H&E-stained kidney sections (n=4-7/group) showing tissue morphology and kidney sections immunostained for apoptotic cell marker, cleaved caspase 3 (CC3) (*n*=5/group) at P28. I and J) Quantification of kidney cortex thickness in the kidney from P28 mice (n=4-7/group) and kidney sections immunostained for apoptotic cell marker, cleaved caspase 3 (CC3) (*n*=5/group) at P28 Statistics: one-way ANOVA followed by the selected pairwise comparisons with Welch’s t-statistics (D, E, I, and J) and Mann-Whitney U test (C). The error bars stand for standard deviation. All data points derive from independent mice.

The kidney is another major affected organ in GRACILE syndrome and in the *Bcs1l^p.S78G^* mice. Proximal tubulopathy with loss of cortex volume is the main histological manifestation (Fig. 3G-J)^4,33,34^. Even though the number of apoptotic cells was not decreased (Fig. 3H and J), the kidney cortex thickness was slightly increased by AAT-*Bcs1l* (Fig. 3G and I), suggesting a systemic growth or metabolic effect from the rescued liver. The treated mice showed normal kidney cortex thickness even at the end-stage, despite increased cell death (Supplementary Fig. 3A-F). The typical albuminuria of the mutant mice^32^ was not prevented by AAT-*Bcs1l* (Supplementary Fig. 3G).

The metabolic status of the *Bcs1l^p.S78G^* mice resembles starvation^4,32^, characterized by hypoglycemia (Fig. 1D) and depleted glycogen stores (Fig. 4A), consistent with earlier reports in both patients and mice with GRACILE syndrome. Another hallmark of the disease is the near-complete absence of white adipose tissue (WAT) depots^4,32^, reflecting reliance on lipids for fuel. Body composition analysis showed that AAT-*Bcs1l* did not correct fat mass at P28 (Supplementary Fig. 4A), possibly because of a persistent need for fuel. Interestingly, by ∼2 months of age (end stage of the survival mice), treated females, but not males, showed normal fat mass (Supplementary Fig. 4A).

**Figure 4.**
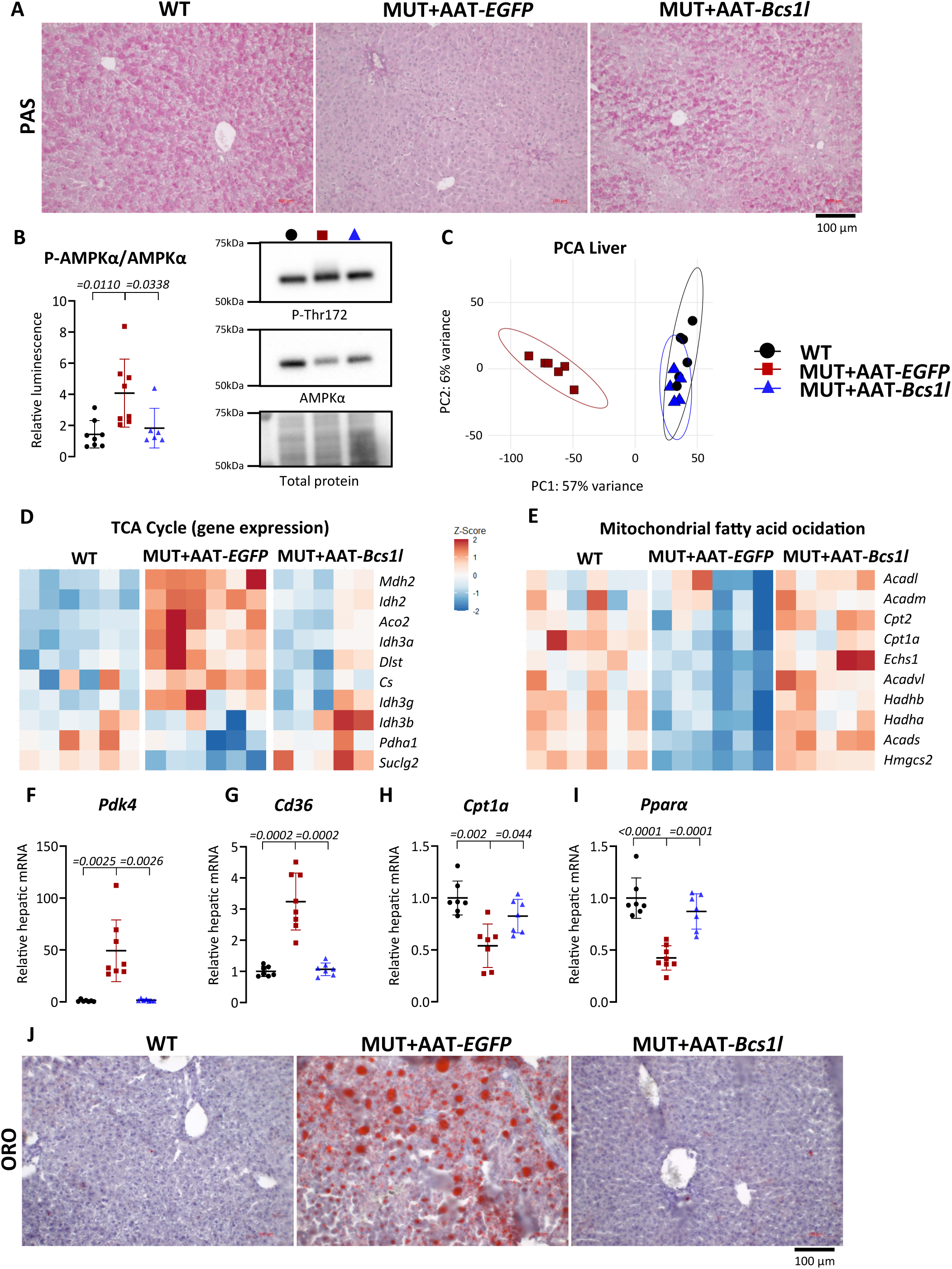
Liver energy metabolism reflects the restored CIII function in hepatocytes. A) Representative images of glycogen stained with periodic acid-Schiff (PAS) on liver sections (n=5/group) from P28 mice. B) Western blot quantification of the phosphorylation status of AMPKα from P28 liver lysates (*n*=6-8/group). C) Principal component analysis (PCA) of differentially expressed genes in the P28 liver transcriptome D and E) Heat map visualization of the top 10 most differentially expressed genes related to TCA cycle and mitochondrial fatty acid oxidation in P28 liver (n=5-6/group). F-I) *Pdk4, Cd36, Cpt1a,* and *Pparα,* mRNA expression from P28 liver (*n*=7-8/group). G) Representative images of Oil-Red-O staining of liver cryosections (n=4/group) showing lipid accumulation at P28. Statistics: One-way ANOVA followed by the selected pairwise comparisons with Welch’s t-statistics (B and F-I). The error bars stand for standard deviation. All data points derive from independent mice. Confidence ellipses indicate 95% confidence intervals for each group (C). The error bars stand for standard deviation. All data points derive from independent mice.

As an established measure of energy deficiency, we examined AMP-dependent protein kinase (AMPK), a central regulator of ATP and glucose availability. While hepatic levels of the AMPK α-subunit was decreased in the mutants^32,34^, its activated form (Thr172-phosphorylated AMPKα) was increased, leading to a higher P-AMPKα/AMPKα ratio (Fig. 4B). This was prevented by AAT-*Bcs1l* (Fig. 4B), indicating improved energy status. Principal component analysis (PCA) of transcriptome data showed a near-complete overlap of liver gene expression profiles between wild-type and AAT-*Bcs1l*–treated mutants, indicating that hepatocyte-specific restoration of CIII function was sufficient to nearly normalize the transcriptome (Fig. 4C and Supplementary Fig. 4B). For example, gene expression changes related to the key energy metabolism pathways TCA cycle and fatty acid oxidation (FAO) were upregulated and downregulated, respectively, in the mutant liver, but not in the AAT-*Bcs1l*-treated livers (Fig. 4D and E). This was further supported by *Pdk4* upregulation (suggesting shutdown of pyruvate dehydrogenase complex by phosphorylation) and *Cd36* upregulation (increased fatty acid uptake from circulation) (Fig. 4F and G), pointing to an attempt to shift from glucose to fatty acid oxidation upon hypoglycemia and glycogen depletion. However, despite the *Cd36* upregulation, expression of the key FAO genes *Cpt1a* (controlling mitochondrial fatty acid import)^36^ and *Pparα* (a major FAO regulator), was markedly decreased (Fig. 4H and I). Together with the microvesicular steatosis (Fig. 4J), these findings suggest a mismatch between fatty acid uptake and FAO capacity in the mutant liver. AAT-*Bcs1l* preserved the glycogen stores and prevented the fat accumulation and basically all energy metabolism-related gene expression changes (Fig. 4A, F–J). These improvements were also reflected in restored expression of *Igf1* and *Ghr*, consistent with improved growth (Supplementary Fig. 4C, D, and Fig. 1E).

The *Bcs1l^p.S78G^* mice show regenerative hepatocyte proliferation, which, in the face of their depleted nucleotide pools and other biosynthetic resources, leads to DNA damage, cell cycle arrest, and senescence^32^. The restoration of CIII function in hepatocytes decreased the DNA damage/senescence marker γH2AX (Fig. 5A) and abolished the cell cycle arrest marker CDKN1A (p21) (Fig. 5B). AAT-*Bcs1l* also fully prevented the hepatic c-MYC upregulation (Fig. 5C), which in both cancer and normal cells can bypass cell cycle checkpoints^37,38^ and the upregulation of the cell cycle markers PCNA and cyclin A2 (Fig. 5D).

**Figure 5.**
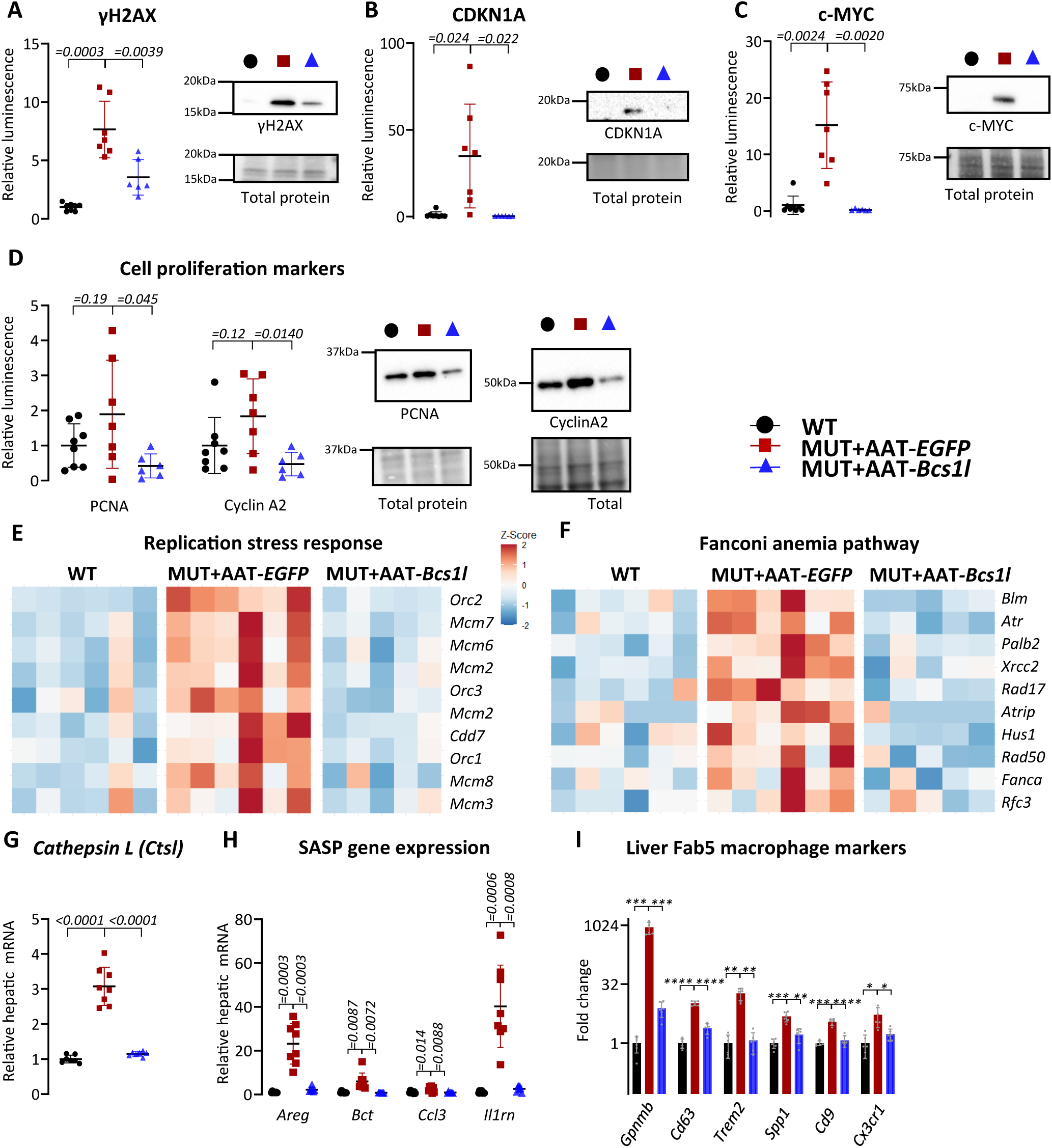
Restoration of CIII function prevents the upregulation of hepatic senescence markers. A-D) Western blot quantification and representative blot of γH2AX, CDKN1A (p21), c-MYC, PCNA, and CyclinA2 from P28 liver lysates (*n*=5-8/group). E and F) Heat map visualization of the top 10 most differentially expressed genes related to replication stress response and Fanconi anemia in P28 liver (n=5-6/group). G) mRNA expression *Cstl* from P28 liver (*n*=7-8/group). H) mRNA expression of senescence-associated secretory phenotype (SASP) genes (*Areg, Bct, Ccl3,* and *Il1rn)* from P28 liver (*n*=7-8/group). I) Fab5 macrophage gene expression in liver transcriptome. Statistics: One-way ANOVA followed by the selected pairwise comparisons with Welch’s t-statistics (A-D and G-I). *\*\*\*\*p ≤ 0.0001, ***p ≤ 0.001, **p ≤ 0.01, *p ≤ 0.05.* The error bars stand for standard deviation. The error bars stand for standard deviation. All data points derive from independent mice.

In line with c-MYC-driven senescence^32^, pathway analyses highlighted activation of replication stress and Fanconi anemia-related genes in the mutant liver. AAT-*Bcs1l* largely prevented these transcriptional changes (Supplementary Fig. 5A and Fig. 5E and F). The upregulation of cathepsin L (*Ctsl*), an aging and senescence marker^39^, was also prevented by AAT-*Bcs1l* (Fig. 5G). Gene expression analyses of senescence-associated secretory phenotype (SASP) showed more than 20-fold increase of the EGF receptor ligands amphiregulin (*Areg*) and betacellulin (*Btc*), and increased expression of several chemokines and cytokines associated with SASP, like chemokine ligand 3 (*Ccl3*) and interleukin 1 receptor agonist (*Il1rn*) in the mutant liver. The expression of all these SASP factors was normal after AAT-*Bcs1l* (Fig. 5H). Markers for a specific fibrogenic bone marrow-derived population of macrophages named Fab5 (CD9^+^TREM2^+^ and expressing SPP1, GPNMB, FABP5, and CD63)^40^ were highly increased in the mutant liver, and this was fully prevented by AAT-*Bcs1l* (Fig. 5I), suggesting that the hepatocyte damage specifically attracts circulating inflammatory cells to the liver and that local correction of the tissue damage prevents this.

### Liver-specific gene therapy improves systemic and skeletal muscle metabolism

Indirect calorimetry using CLAMS equipment revealed a significant alteration in whole-body substrate utilization in the mutant mice (Fig. 6A). Consistent with our previous findings^28,33^, their respiratory exchange ratio (RER) of <0.9 indicated a shift from glucose to fatty acid oxidation, consistent with the hypoglycemia and glycogen depletion. AAT-*Bcs1l-*treated mutants had significantly higher RER values, demonstrating a systemic preservation of glucose utilization and energy metabolism (Fig. 6A). In line with the improved systemic energy metabolism, AAT-*Bcs1l* treatment strikingly shifted transcriptome-wide expression changes in the skeletal muscle towards normal (Fig. 6B). qPCR for total and virally expressed *Bcs1l* confirmed that AAT-*Bcs1l* did not give non-specific expression in the skeletal muscle (Supplementary Fig. 6A), and the effect originated in the liver. Upon AAT-*Bcs1l*, the expression of ∼96% of the dysregulated genes in the mutant skeletal muscle was significantly shifted towards WT levels (Fig. 6C, Supplementary Fig. 6B). Pathway analyses highlighted the upregulation of energy metabolism-related processes in the mutant skeletal muscle (Supplementary Fig. 6C and Fig. 6D-F). Similar to the liver, the mutant skeletal muscle showed upregulation of *Ppargc1a (Pgc-1α)*, which was effectively prevented by AAT-*Bcs1l* (Fig. 6G). In response to decreased glucose availability, skeletal muscle also compensates by upregulating FAO. Consistent with this, the expression of *Pdk4* and *Cd36* was markedly elevated in the mutant muscle (Fig. 6H and I), indicating a metabolic shift from glucose to fatty acid utilization. AAT-*Bcs1l* significantly attenuated the *Pdk4* and *Cd36* upregulation (Fig. 6H and I). In summary, restoring CIII function and mitochondrial respiration in the liver led to robust correction of markers for systemic and skeletal muscle energy metabolism.

**Figure 6.**
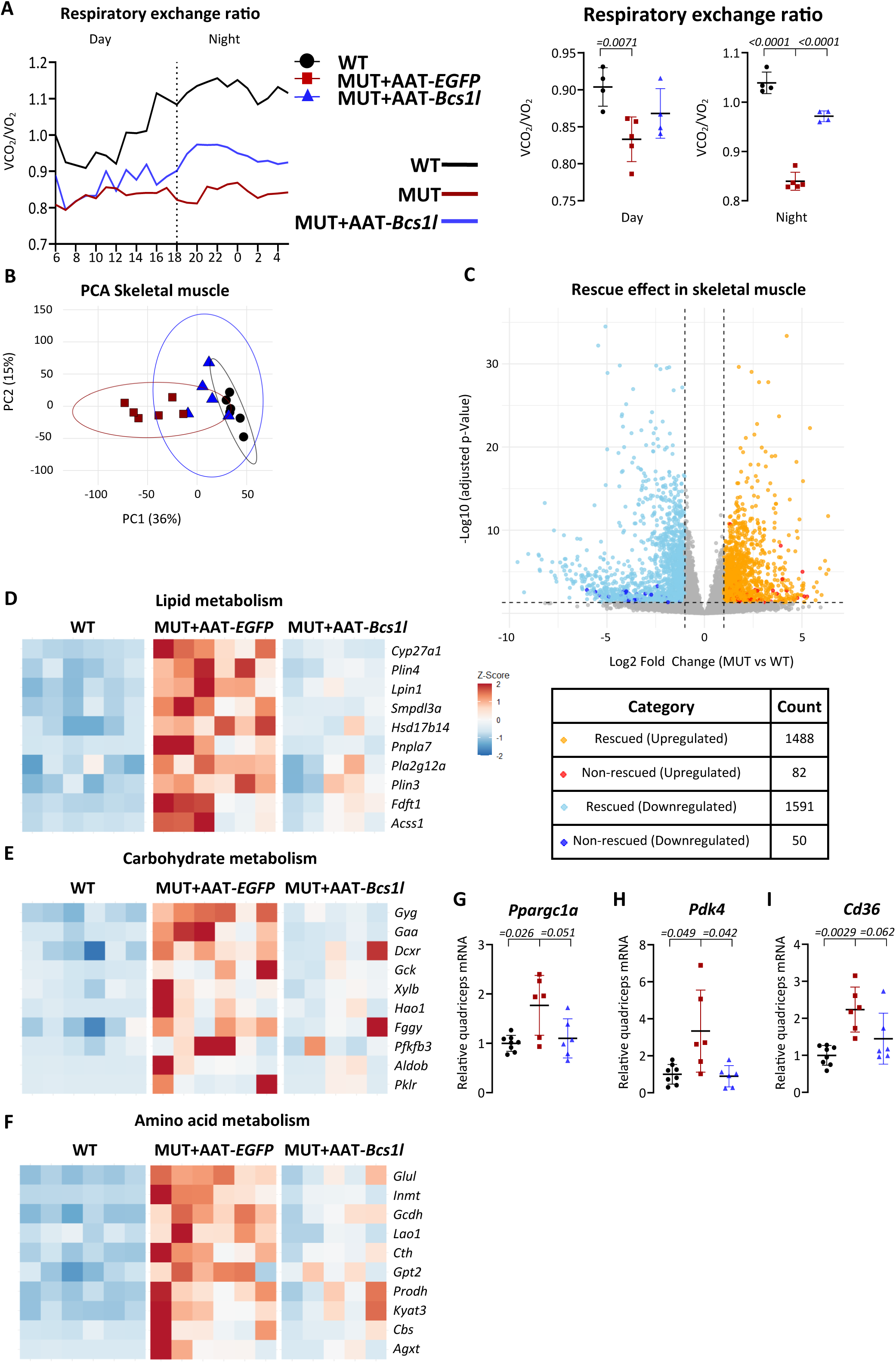
Liver-targeted gene replacement improves skeletal muscle metabolism. A) A Comprehensive Laboratory Animal Monitoring System (CLAMS) apparatus was used to measure O_2_ consumption and CO_2_ production at age P26-27. Circadian rhythm of the respiratory exchange ratio and the average 12-h respiratory exchange ratio of daytime and nighttime measurements are shown. B) PCA of differentially expressed genes in the P28 skeletal muscle transcriptome. C) Rescued genes shown in the volcano plot are transcripts whose expression levels were restored towards wild type, while non-rescued genes remained dysregulated. All other genes are shown in grey. D-F) Heat map visualization of the top 10 most differentially expressed genes related to lipid, carbohydrate, and amino acid metabolism in P28 skeletal muscle (n=5-6/group). G-I) *Ppargc1*, *Pdk4* and *Cd36* mRNA from P28 skeletal muscle (n=6-8/group). Statistics: One-way ANOVA followed by the selected pairwise comparisons with Welch’s t-statistics (G-I). The error bars stand for standard deviation. All data points derive from independent mice. Confidence ellipses indicate 95% confidence intervals for each group (B).

## Discussion

Despite the rapidly expanding knowledge about molecular disease mechanisms, for most genetic diseases, such as mitochondrial diseases, gene therapy will likely remain the only effective treatment strategy. Here, we present the first preclinical gene therapy trial in a mouse model of a human CIII deficiency, using a robust model based on the GRACILE syndrome patient mutation^4,5^. In contrast to the patients, who are very sick already at birth^4^, the mice are healthy until weaning age^28,33,34^, providing a useful presymptomatic window. We show that both broad (CAG promoter) and liver-specific (alpha1-antitrypsin promoter) rAAV-based gene replacement in the juvenile mice prevented the early metabolic crisis and doubled the survival from one month to nearly two months. AAT-*Bcs1l* prevented the liver disease typical of GRACILE syndrome and, in a milder form, of the mouse model^4,34,41^, and abrogated hepatocyte senescence and hepatic inflammation. These results support the view that mitochondrial dysfunction in hepatocytes underlies the liver disease and that the liver disease plays an important role in the compromised systemic energy metabolism. For example, the growth of the AAT-*Bcs1l*-treated mutant mice was improved, which was likely due to their improved energy metabolism. Interestingly, the poor growth of the *Bcs1l^p.S78G^* mice may also be a part of their progeroid phenotype, which includes osteopenia, kyphosis, and loss of white adipose tissue^32^.

During fasting, but also in the chronic starvation-like metabolic state in mitochondrial diseases, fatty acids are mobilized from adipose tissue to the liver^38^. Despite apparently normal food intake, the *Bcs1l^p.S78G^* mice face hypoglycemia and glycogen depletion, forcing a shift from glucose to fatty acid utilization^8,34^. To facilitate this, the mutant hepatocytes upregulate fatty acid uptake, but persistent micro vesicular fat accumulation suggests insufficient hepatic FAO, a heavily mitochondria-dependent catabolic process. Restoring hepatic mitochondrial respiration with AAV-*Bcs1l* reversed the markers for the glycolysis-to-FAO metabolic switch and prevented the glycogen depletion and hypoglycemia. Indirect calorimetry data also reflected the prevention of glycolysis-to-FAO metabolic switch at the whole-body level. As an example of extrahepatic effects potentially mediated by the improved systemic energy metabolism, the liver-specific gene therapy improved kidney cortex mass, which is decreased by proximal tubulopathy in both patients and the mutant mice^4,33,34^. Because the skeletal muscle shows ∼75% loss of CIII activity in the juvenile *Bcs1l^p.S78G^* mice^28^ and is a large energy-consuming organ, we wanted to assess its potential role in the gene therapy effects. Quite astonishingly, AAT-*Bcs1l* robustly prevented the transcriptome-wide alterations in the skeletal muscle, underscoring a strong metabolic effect from the liver. Notably, the liver-specific gene therapy prevented the glycolysis-to-FAO switch also in the skeletal muscle. Together, these findings demonstrate that restoring hepatic CIII function exerts systemic benefits on peripheral tissues, improving overall energy homeostasis.

The gene therapy led to robust prevention of liver disease and metabolic alterations in our mouse model, yet the survival still decreased sharply at ∼2 months. Genomic integration did not further extend the survival, indicating that the eventual deterioration was driven primarily by CIII deficiency in non-hepatic tissues. Intraperitoneal injection of rAAVs with broader CAG promoter-driven expression provided only a modest additional survival benefit. Therefore, less efficient biodistribution in harder to transduce tissues like skeletal muscle likely limited the rescue effect. Optimizing delivery strategy and/or capsid may improve the biodistribution. Loss of episomal expression over time may also be a challenge, particularly in patients with much longer life span than mice. On the other hand, recent encouraging result from CI- and CIV-related encephalopathy mouse models showed long-lasting (15 moths) effects with rAAV-based gene replacement in the brain^42^. Repeated rAAV administration is hindered by immune responses against AAV capsids^43,44^. Therefore, alternative delivery systems such as lipid nanoparticles (LNPs)^45^, engineered virus-like particles (eVLPs)^46^, and extracellular vesicle–derived vectors (eDVs)^47^ are emerging as promising tools that can bypass immunogenicity, enable repeated dosing, and potentially broaden tissue tropism. Finally, with the advent of CRISPR-based genome editing, including base editing^48^ and prime editing^49^, it is becoming increasingly feasible to achieve permanent correction of disease mutations, potentially overcoming the durability limitations of conventional rAAV-based gene replacement. The GRACILE syndrome mutation is difficult to correct using base editing (our unpublished data from patient fibroblasts), but prime editing strategies should work and can be designed for further preclinical trials in the *Bcs1l^p.S78G^* mouse model.

In summary, our results demonstrate the first successful preclinical gene therapy trial in a mouse model of CIII deficiency. The most striking outcome of this study was the degree of systemic rescue achieved through hepatocyte-specific restoration of BCS1L and CII function. The results are potentially translatable to mitochondrial disease patients with *BCS1L* mutations, most of which cause clearly milder phenotypes than the GRACILE syndrome mutation^50^. The work also sheds light on the role of the liver in the systemic manifestations, such as hypoglycemia and loss of adipose tissue, in CIII deficiency. These results provide further proof that targeting a single metabolically dominant organ can delay systemic disease in mitochondrial disorders. Liver-directed gene therapy may therefore represent a promising strategy for multiorgan mitochondrial diseases with prominent hepatopathy.

## Materials and Methods

### Cloning and virus production

Mouse *Bcs1l* coding sequence (*MmBcs1l*) was PCR-amplified from mouse tissue cDNA (primers EcoRI-MmBcs1l: 5’-ATGAATTCACCATGCCATTTTCAGACTTTGTTCTG-3’ and MmBcs1l-STP-HindIII: 5’-ATAAGCTTTCACCTCAGAGATTCAATGTTGT-3’). The insert was ligated into pBluescript, sequenced, and then subcloned into a standard rAAV production vector pSUB-CAG-WPRE for broad expression under the chicken β-actin promoter and the CMV immediate early enhancer, facilitating ubiquitous gene expression across cell types (RRID: Addgene_119227), and into pAAV2-LSP1-PB(TR)-EGFP vector^29^. The latter allows hepatocyte-specific expression under the human ApoE enhancer and α1-antitrypsin promoter (generously provided by Professor Ian Alexander, University of Sydney). The parental vector encoding enhanced green fluorescent protein (EGFP) was used as a control. Along with the expression cassette, the vectors also contained flanking PiggyBac transposon recognition sequences, which allow for genomic integration upon parallel expression of the PiggyBac transposase^29^. Serotype 9 viral particles were produced by the AAV Gene Transfer and Cell Therapy Core Facility of the University of Helsinki.

### Mouse breeding and husbandry

The animal facilities of the University of Helsinki maintained the mice on the C57BL/6JCrl background (Harlan stock 000016). As the GRACILE syndrome is a classic recessive disease^5^, the patients nor the mouse model has a heterozygous phenotype^6^. Hence, *Bcs1l* wildtype or heterozygous animals were used as healthy controls (wild type, WT, by phenotype). Both males and females were used, and data are shown separately for them only if a significant sex difference was observed or known.

The mice were housed in individually ventilated cages with a 12-hour light/12-hour dark cycle at a temperature of 22-23°C, and they had ad libitum access to water and food (2018 Teklad global 18% protein rodent diet, Envigo). Mouse health was monitored by manual behavioral scoring and weighing according to the ethical permit. Samples were collected on postnatal day 28 (P28) or according to the survival of the mice. In survival analysis, the mice were euthanized when weight loss was greater than 15% of the maximum weight of the individual mouse.

The animal studies were approved by the animal ethics committee of the State Provincial Office of Southern Finland (ESAVI/16278/2020 and ESAVI/31141/2023) and were performed according to FELASA (Federation of Laboratory Animal Science Associations) guidelines. The animal work and experimental setup were designed following 3R principles.

### rAAV administration

Presymptomatic (postnatal day 19-23, P19-23) *Bcs1l^p.S78G^*; *mt-Cyb^p.D254N^* mice were injected intraperitoneally with 100 µl saline containing 5×10^10^ viral particles encoding EGFP or wild-type *Bcs1l*. rAAVs encoding EGFP were used as a control for any vector-related effects, as is common in preclinical gene therapy trials. However, because the EGFP-expressing mutant mice were comparable to the non-injected mutant mice, we later omitted the AAV-EGFP injections and used non-injected homozygotes as controls. We determined the dosing using i.p. injection and by testing doses of 1×10^10^, 5×10^10^ and 25×10^10^ viral particles expressing EGFP in wild-type mice. We selected the 5×10^10^ dose based on the visual EGFP expression in the transduced liver.

### Assessment of body composition

The echoMRI-based MiniSpec Body Composition Analyzer (Bruker, USA) was used to quantify fat mass.

### Sample collection

The mice were euthanized by cervical dislocation. Before sample collection, the mice underwent a short 2-hour fasting period to avoid exacerbation of hypoglycemia in the mutant mice, and all samples were obtained at a similar time of the day, during the light period. Blood glucose was measured with a quick meter (Freestyle Lite, Abbott, USA) from the blood within the body cavity while collecting the other samples. Tissues were either placed in 10% histology-grade formalin or snap-frozen in liquid nitrogen and stored at −80°C.

### Quantitative PCR (qPCR) and RNA sequencing

Total RNA was extracted from the snap-frozen tissue samples with RNAzol RT reagent (Sigma-Aldrich). qPCR was performed from the cDNA using EvaGreen- and Phire II Hot Start DNA polymerase-based detection chemistry^28^. CFX96 thermocycler and CFX Manager software (Bio-Rad) were utilized to perform the qPCR and data analysis. LinRegPCR software^51^ was used to calculate PCR efficiency. All primers used can be found in Supplementary Table 1. *Gak* and *Rab11a* served as reference genes.

RNA sequencing and primary bioinformatics analysis were performed by Novogene Inc. (Beijing). Downstream analyses were conducted in R (version 4.4.2) using the DESeq2 package^52^. Variance stabilizing transformation (VST)-normalized read counts generated with DESeq2 were used for principal component analysis (PCA). Differential expression analysis was performed with DESeq2, and genes were considered significantly differentially expressed at an adjusted p-value < 0.05 and an absolute log2 fold change > 1. Pathway analysis was conducted using Gene Set Enrichment Analysis (version 4.3.3)^53^. Data visualization, including bubble plots, PCA plots, and heatmaps, was performed in RStudio. In the heatmaps, top 10 differentially expressed genes are shown for each pathway or gene ontology class.

### SDS-PAGE, blue-native PAGE, and Western blot

For Western blot analyses of tissue processing, SDS-PAGE analysis, and immunoblotting were performed as described in the protocol^32^.

For blue native gel electrophoresis (BNGE), liver mitochondria were first freshly isolated using the previously described protocol^28^. Then they were solubilized by adding 6 mg digitonin per mg of protein in a cold buffer comprising (50 mM Bis-Tris-Cl^+^, 50 mM NaCl, 1.4 mg/ml digitonin, 10% glycerol, 1 mM EDTA, protease inhibitor mix, pH of 7.0). The sample lysates were clarified by centrifugation at 18,000g at +4°C for 6 minutes. Coomassie Blue G-250 (1.5 mg/ml) was added to the supernatants, and 10μg solubilized mitochondrial protein was separated using 3–12% NativePAGE^TM^ Bis-Tris gradient gels (Invitrogen) and electroransferred onto PVDF filters as described^22^.

For the analysis of albumin and major urinary proteins (MUPs), urine samples were heated and mixed with Laemmli sample buffer (LSB), and an equivalent of 1 µl of urine was loaded onto 4– 20% tris-glycine polyacrylamide gel electrophoresis (PAGE) gels. The gels were stained with Coomassie G-250, and images were captured using a flatbed scanner. All the samples were randomized before running and processing for quantification. For representative blots, individual samples were pooled from each experimental group to obtain an average signal, which represents each group.

### Assessment of respiratory chain enzymatic activities and quantification of ATP

CIII activity was measured using a spectrophotometric method that involved monitoring the reduction of cytochrome c, sensitive to antimycin A and myxothiazol, with decylubiquinol serving as the electron donor ^28^. The CIII activity data were normalized to protein content.

For the enzymatic quantification of ATP from the liver, we used a published protocol^54^.

### Tissue histology and immunohistochemistry

Formalin-fixed paraffin-embedded tissues underwent standard procedures for general histological assessment, including hematoxylin and eosin (H&E) staining and glycogen detection using periodic acid-Schiff (PAS) staining. Additionally, frozen liver sections fixed in formalin and saturated with 30% w/v sucrose were subjected to the standard Oil Red O (ORO) staining to detect triglycerides. The antigen retrieval of paraffin sections for cleaved caspase 3 and HSP60 staining (supplementary Table 2) was done by immersing the slides in 10 mM Tris-Cl, pH 9.0; 1mM EDTA, and boiling for 15 minutes. After incubating the sections in the primary antibody, ImmPress peroxidase- or alkaline phosphatase polymer-conjugated of secondary antibodies (Vector Laboratories Inc) were added. Nitroblue tetrazolium was used to visualize alkaline phosphatase and diaminobenzidine peroxidase activity, respectively. Nuclear Fast Red (Sigma-Aldrich) and hematoxylin were used as nuclear counterstains for alkaline phosphatase and peroxidase stainings, respectively.

### Statistics

All samples were randomized before quantification and processing. Group differences were assessed using Welch’s t-test, or the Mann–Whitney U test when appropriate, for comparisons between two groups. For comparisons among multiple groups, one-way ANOVA followed by pre-selected pairwise comparisons using Welch’s t-statistics was applied. All pairwise comparisons were conducted using two-sided tests. Survival analyses were performed using the log-rank (Mantel–Cox) test. Statistical analyses were conducted with GraphPad Prism version 10 (GraphPad Software Inc.). Unless otherwise specified, error bars in figures represent the mean ± 95% confidence interval (CI). A p-value < 0.05 was considered statistically significant. Statistical tests for RNA-seq data were done with R Studio. PCA outliers were identified using the Mahalanobis distance method and consecutively removed from further analysis.

### Limitations of the study

While AAT-*Bcs1l*-mediated hepatic rescue alleviated systemic metabolic stress and improved transcriptome signatures of skeletal muscle, we did not perform functional assays of muscle performance, and thus, the physiological impact of the rescue remains unknown. As rAAV-mediated expression is episomal and declines over time, this needs to be taken into account when considering translation to humans with much longer lifespans. Gradual failure of the kidney was the most likely explanation for the eventual deterioration of the treated mice, but it was not possible to prove this. Finally, while the transcriptomic analyses provided great mechanistic insight, they did not allow conclusions about protein- and metabolite-level effects of the gene therapy.

### Data availability

Any dataset published here is available from the corresponding author upon request.

## Acknowledgements

We thank Vilma Wanne, Oliver Ros, Divya Upadhyay, and Katariina Kemppainen for technical assistance, and Professor Ian Alexander (University of Sydney) for providing the pAAV-LSP1 plasmids. We thank the core facilities of the University of Helsinki: FIMM Digital Microscopy and Molecular Pathology Unit and the Finnish Centre for Laboratory Animal Pathology (Faculty of Veterinary Medicine) for processing histological samples, Biomedicum Imaging Unit for microscopy services, and the Laboratory Animal Center of the University of Helsinki for the animal husbandry. We acknowledge the funding from Samfundet Folkhälsan, Jane and Aatos Erkko Foundation, Sigrid Juselius Foundation, University of Helsinki, The Foundation for Pediatric Research, Finska Läkaresällskapet, Medicinska Understödsföreningen Liv och Hälsa, Magnus Ehrnrooth Foundation, The Finnish Academy of Science and Letters, and Finnish Doctoral Programme in Oral Sciences (FINDOS), American and European Society of Gene & Cell Therapy (ASGCT and ESGCT) (travel grants to RB). Graphical illustrations were created with BioRender (https://www.biorender.com/). ChatGPT is acknowledged for language editing and R programming language assistance.

## Author contributions

R.B., J.P., V.F., and J.K. designed the study. R.B. wrote the first manuscript draft and prepared the figure panels. R.B., J.P., C.K., and J.K. performed the animal experiments and sample collection. R.B., J.P., and J.K. performed the histological analyses. The contributions to the other methods were the following: body composition analyses (R.B. and J.P.), SDS-PAGE and Western blot analyses (R.B., N.S., and J.K.), Blue-Native PAGE (R.B. and J.P.), qPCR (R.B., N.S.), immunohistochemistry (J.K.), and ATP measurements (R.B. and J.P.), transcriptomics (R.B. and C.K.). R.B. was responsible for the statistics. R.B., V.F., and J.K. acquired funding for the project. All authors critically read and commented on the manuscript, and R.B. and J.K. revised it accordingly.

## Figure legends

**Supplementary Figure 1.**
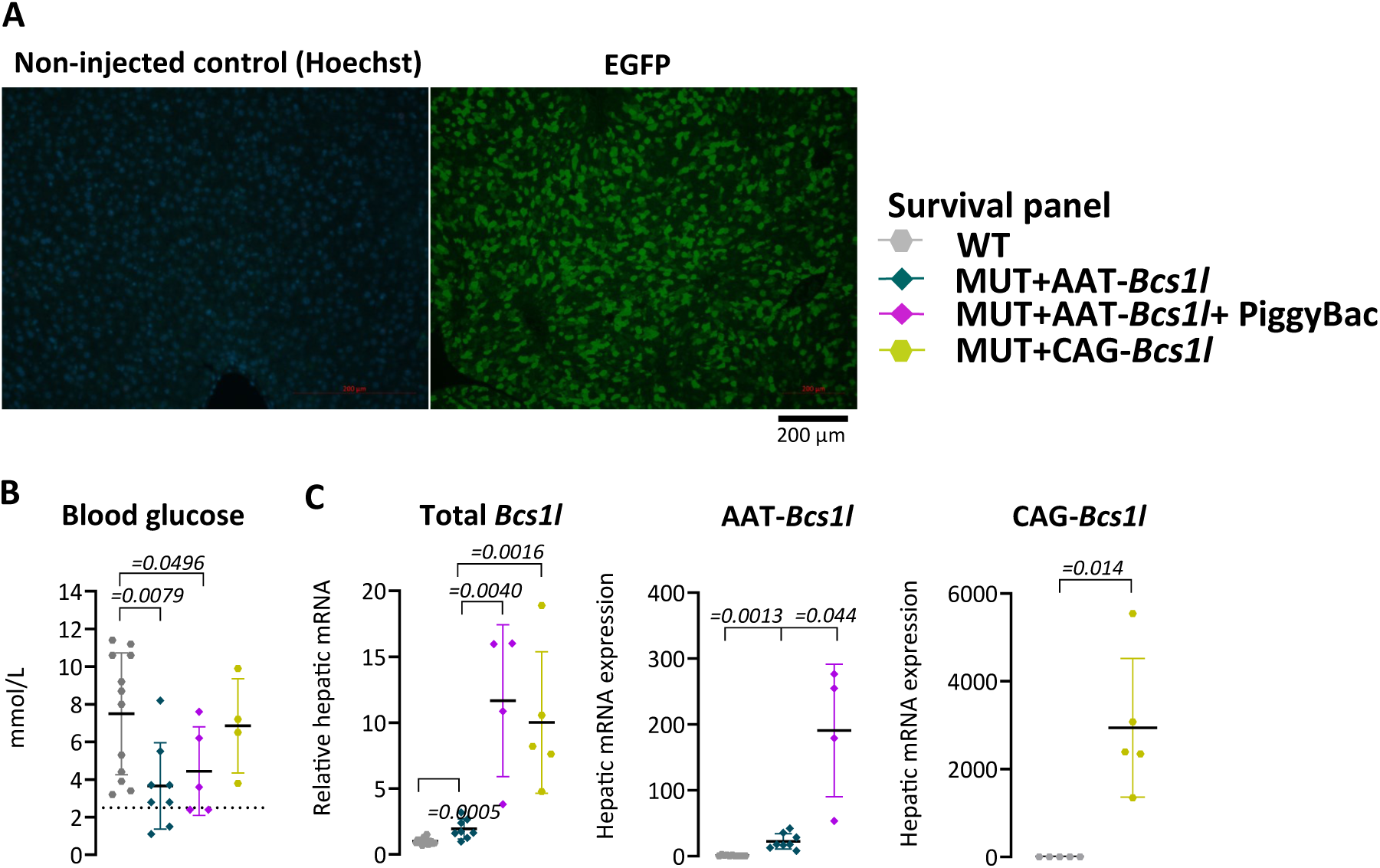
Transduction efficiency of the liver by rAAVs. A) Representative transduction efficiency with the hepatocyte-targeted viral vector. B) Blood glucose levels at the end stage of the survival panels (*n*=4-12/group). The dotted line indicates the critical level of glucose (<2.5mmol/L) predicting spontaneous death. C) Total and virally expressed *Bcs1l* mRNA at the end stage of the survival analysis (*n*=4-17/group). Statistics: One-way ANOVA followed by the selected pairwise comparisons with Welch’s t-statistics (B). Mann-Whitney U test (C). The error bars stand for standard deviation. All data points derive from independent mice.

**Supplementary figure 2.**
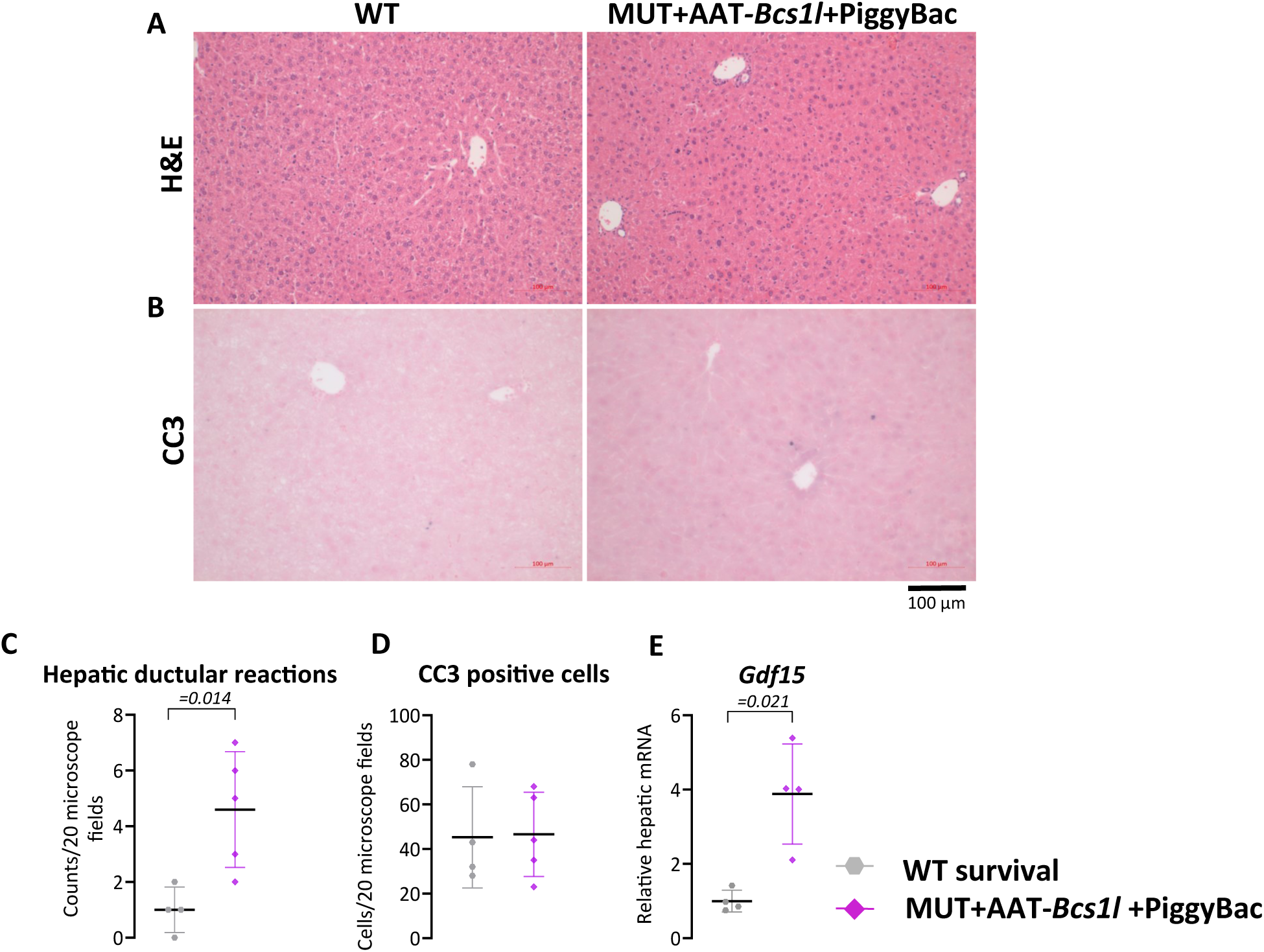
End-stage liver in mutant mice after hepatocyte-targeted gene replacement. A and B) Representative images of H&E-stained liver sections (*n*=5/group) showing tissue. morphology and immunostainings of liver sections for apoptotic cell marker, CC3 (*n*=5/group) at the end stage of the survival analysis. C and D) Quantification of the hepatic ductular reactions and the apoptotic cells from H&E-stained sections and CC3-immunostained sections, respectively. E) *Gdf15* mRNA expression from livers of the survival groups (*n*=4/group). Statistics: Welch’s *t*-test. The error bars stand for standard deviation. All data points derive from independent mice.

**Supplementary figure 3.**
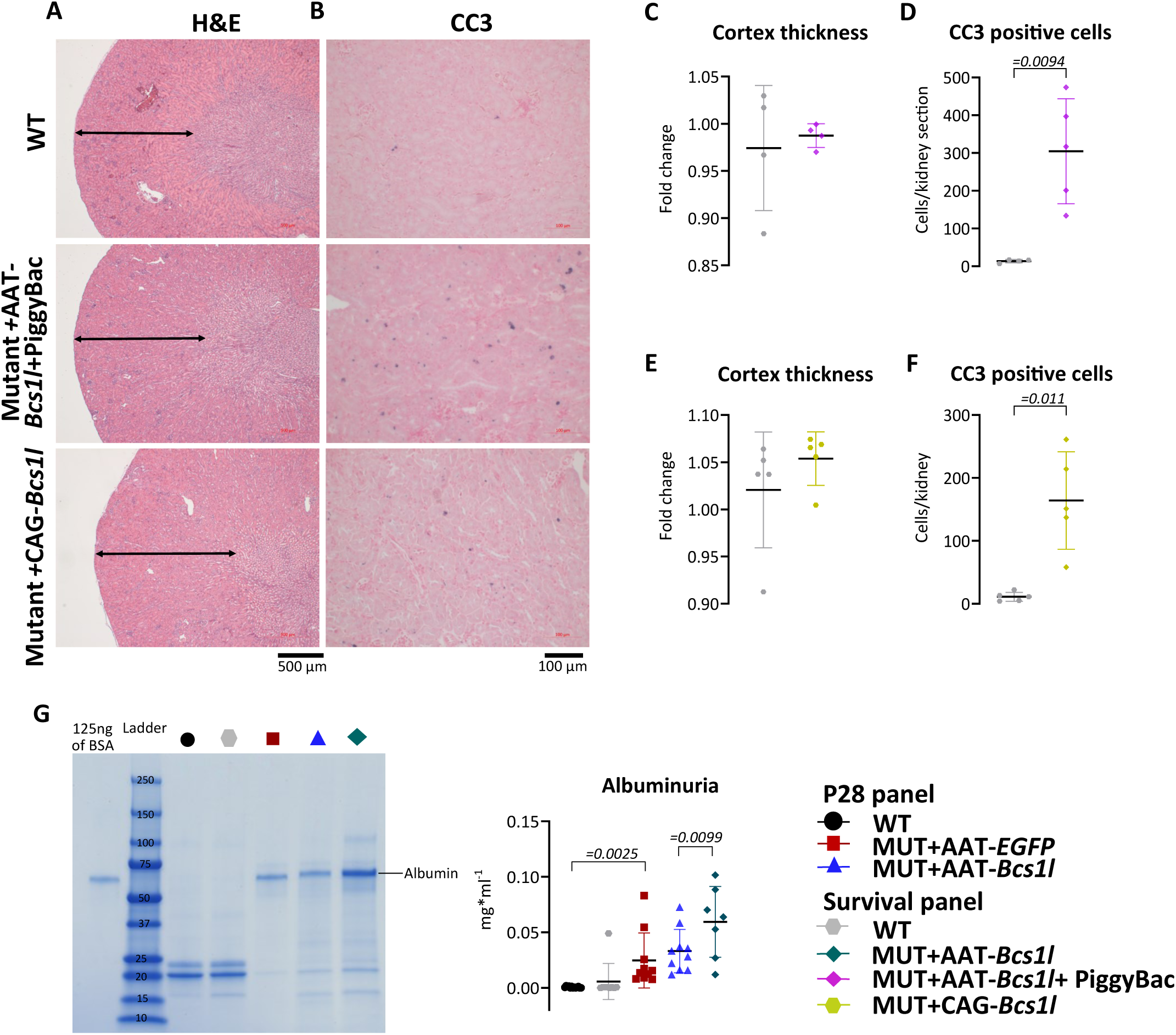
Effect of hepatocyte-targeted gene replacement on kidney disease. A and B) Representative images of H&E-stained kidney sections (n=4-7/group) showing tissue morphology and immunostainings of CC3 (n=5/group) showing apoptotic cells in the kidney, respectively, from survival AAT-*Bcs1l*+PiggyBac- and CAG-*Bcs1l*-treated mice. C-F) Quantification of kidney cortex thickness and apoptotic cells from H&E-stained and CC3- immunostained sections, respectively, from survival AAT-*Bcs1l*+PiggyBac- and CAG-*Bcs1l*-treated mice G) Albumin level in urine from P28, and survival of AAT-Bcs1l-treated mice. Statistics: One-way ANOVA followed by the selected pairwise comparisons with Welch’s t-statistics (G) and Welch’s *t*-test (C-F). The error bars stand for standard deviation. All data points derive from independent mice.

**Supplementary figure 4.**
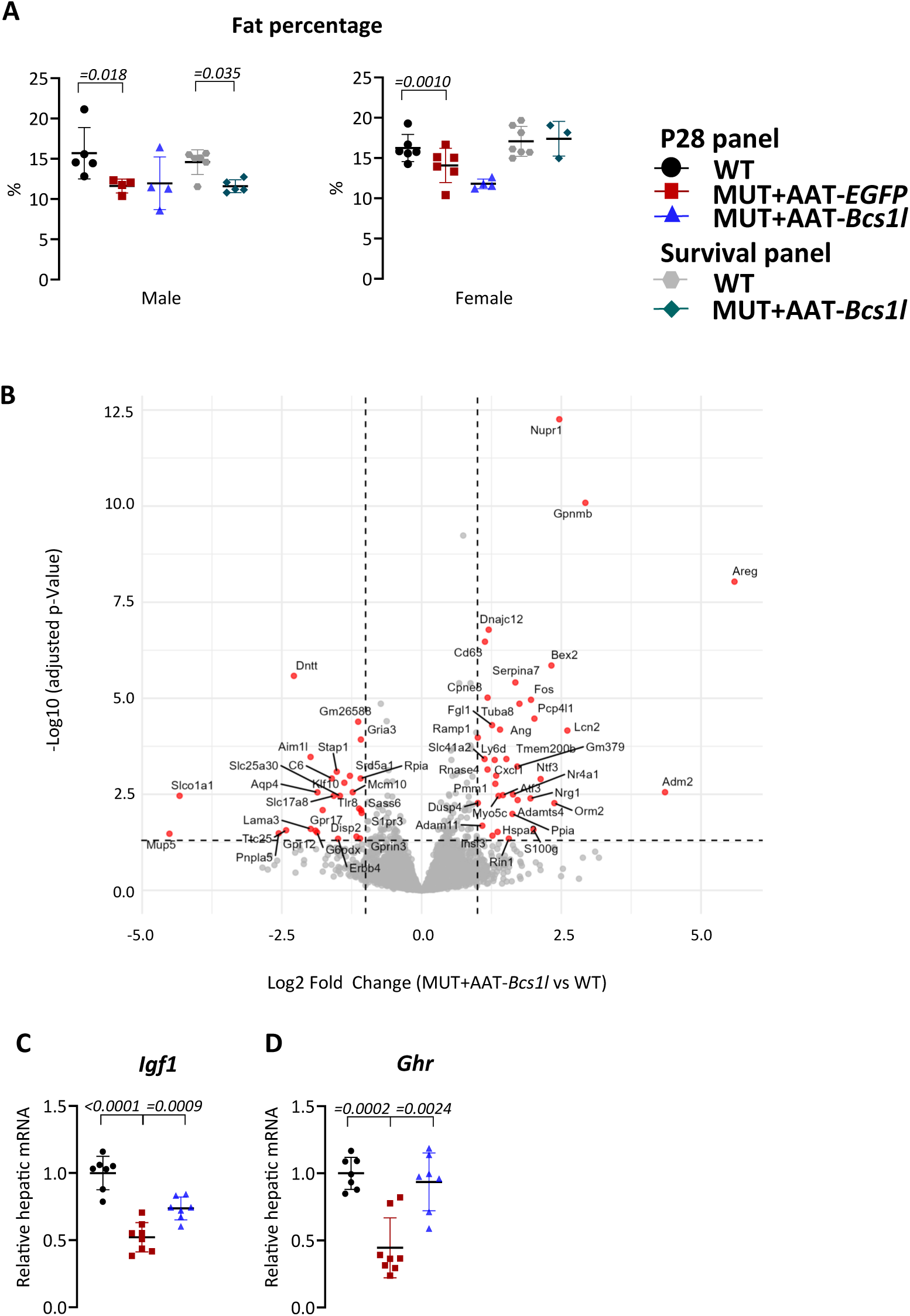
Effect of hepatocyte-targeted gene replacement on hepatic energy metabolism. A) Echo-MRI quantification of body fat percentage in P28, and survival AAT-*Bcs1l*-treated mice. B) Volcano plot showing differentially expressed genes in the liver between AAT-*Bcs1l*-treated mutants and WT mice. C) Hepatic mRNA expression of insulin-like growth factor 1 (*Igf1*), and growth hormone receptor (*Ghr*) at P28 (*n*=7-8/group) Statistics: One-way ANOVA followed by the selected pairwise comparisons with Welch’s t-statistics (A, C and D). The error bars stand for standard deviation. All data points derive from independent mice.

**Supplementary figure 5.**
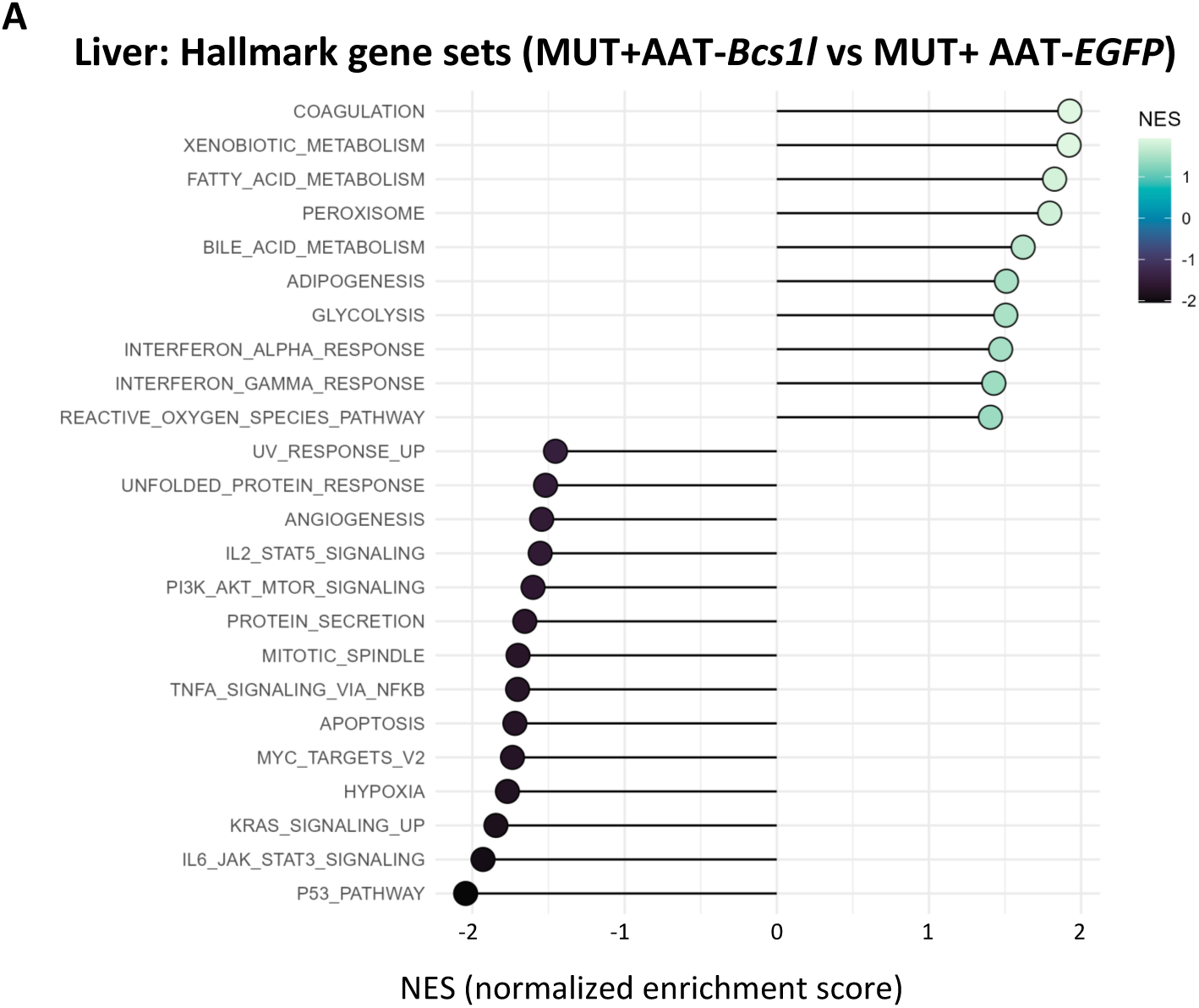
Effect of hepatocyte-targeted gene replacement on different cellular pathways. A) Differentially regulated pathways in the P28 mutant liver transcriptome.

**Supplementary figure 6.**
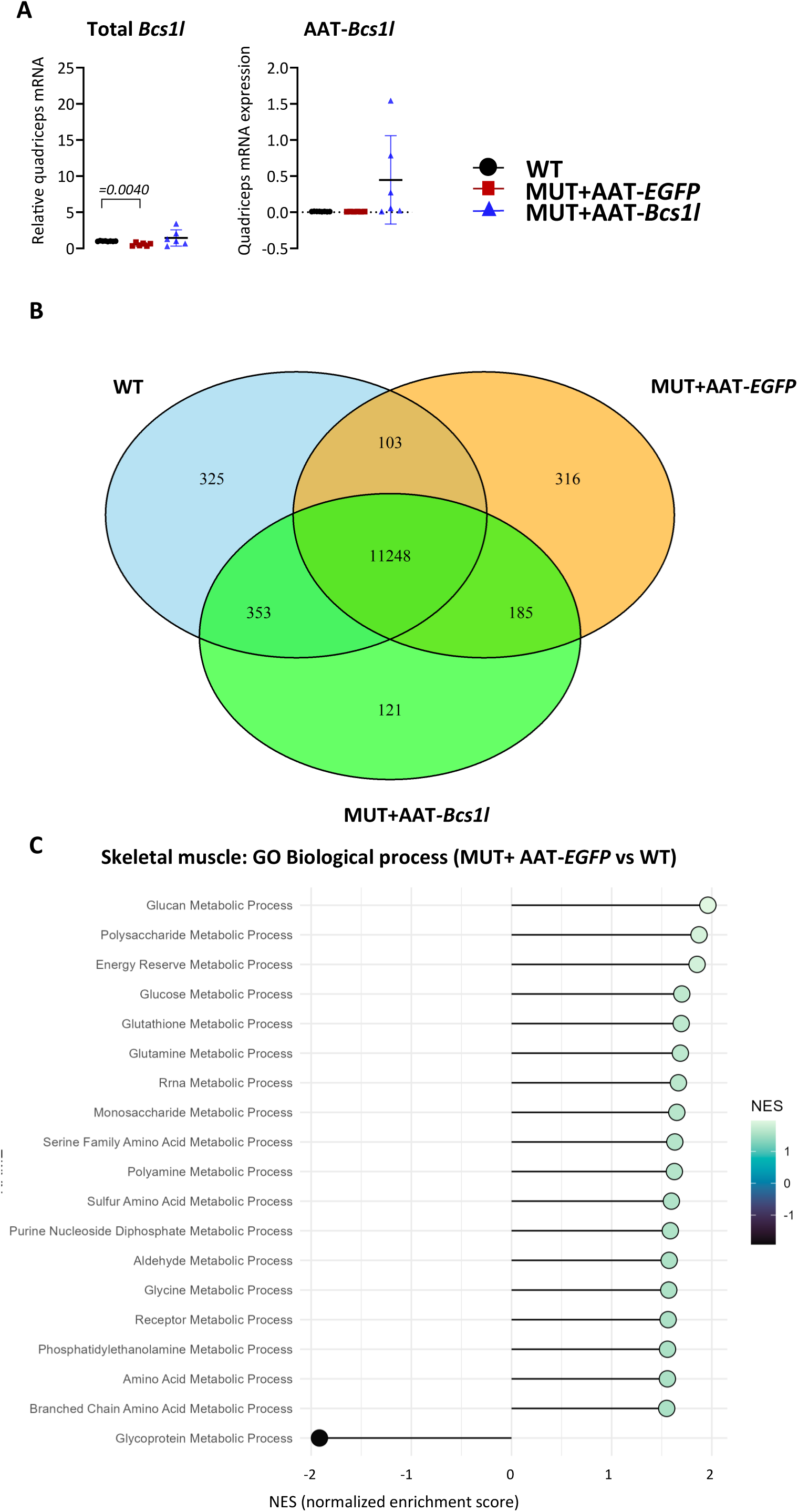
Effect of hepatocyte-targeted gene replacement on muscle energy metabolism. A) Virally expressed and total *Bcs1l* mRNA at P28 quadriceps (*n*=6/group). B) Venn diagram showing differentially expressed genes in P28 skeletal muscle between groups after hepatocyte-targeted gene replacement therapy. C) Differentially regulated pathways in the P28 mutant skeletal muscle transcriptome. Statistics: Mann-Whitney U test (A). The error bars stand for standard deviation. All data points derive from independent mice.

**Supplementary Table 1:**
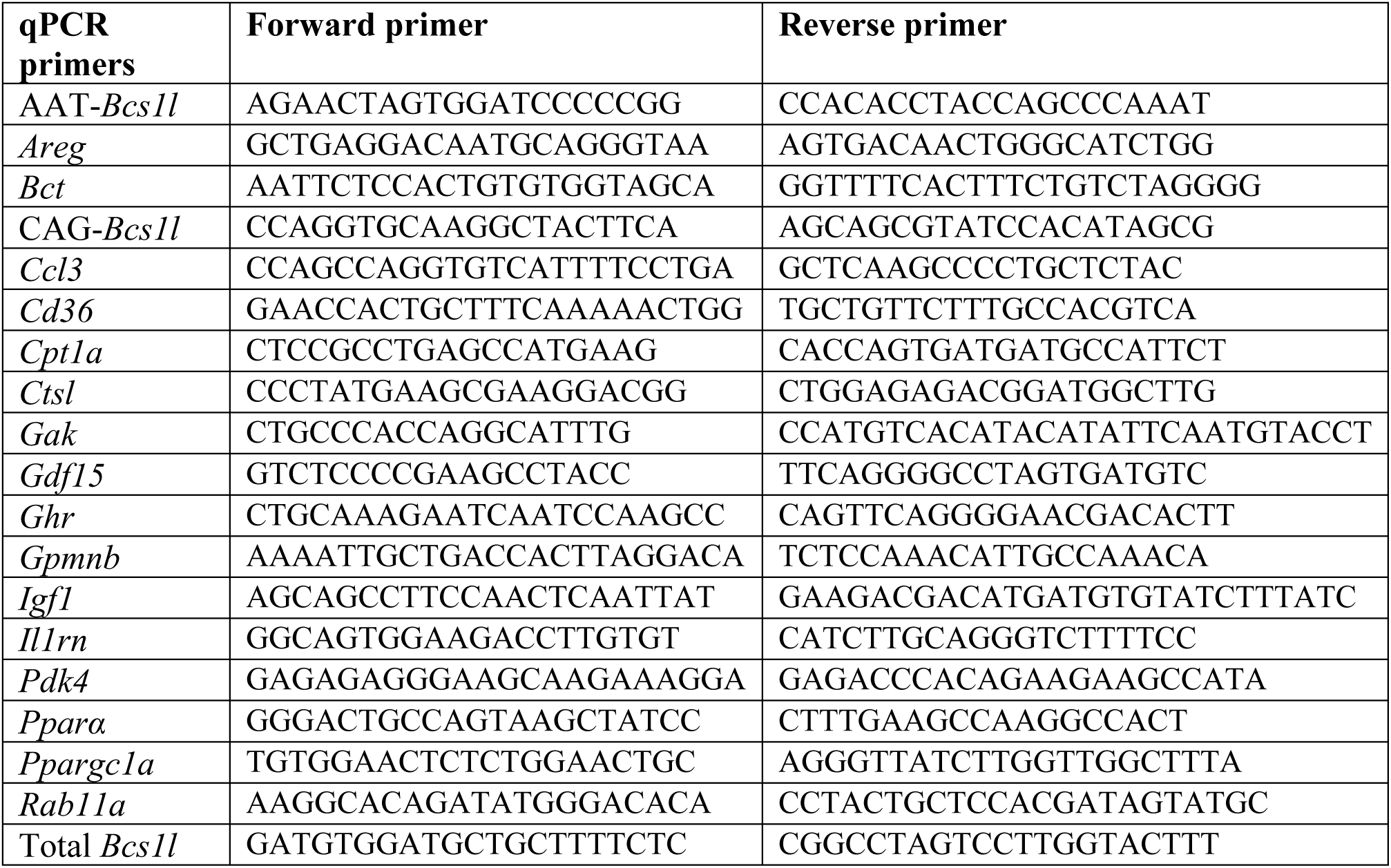
Oligonucleotide sequences in 5’ to 3’ direction.

**Supplementary Table 2:**
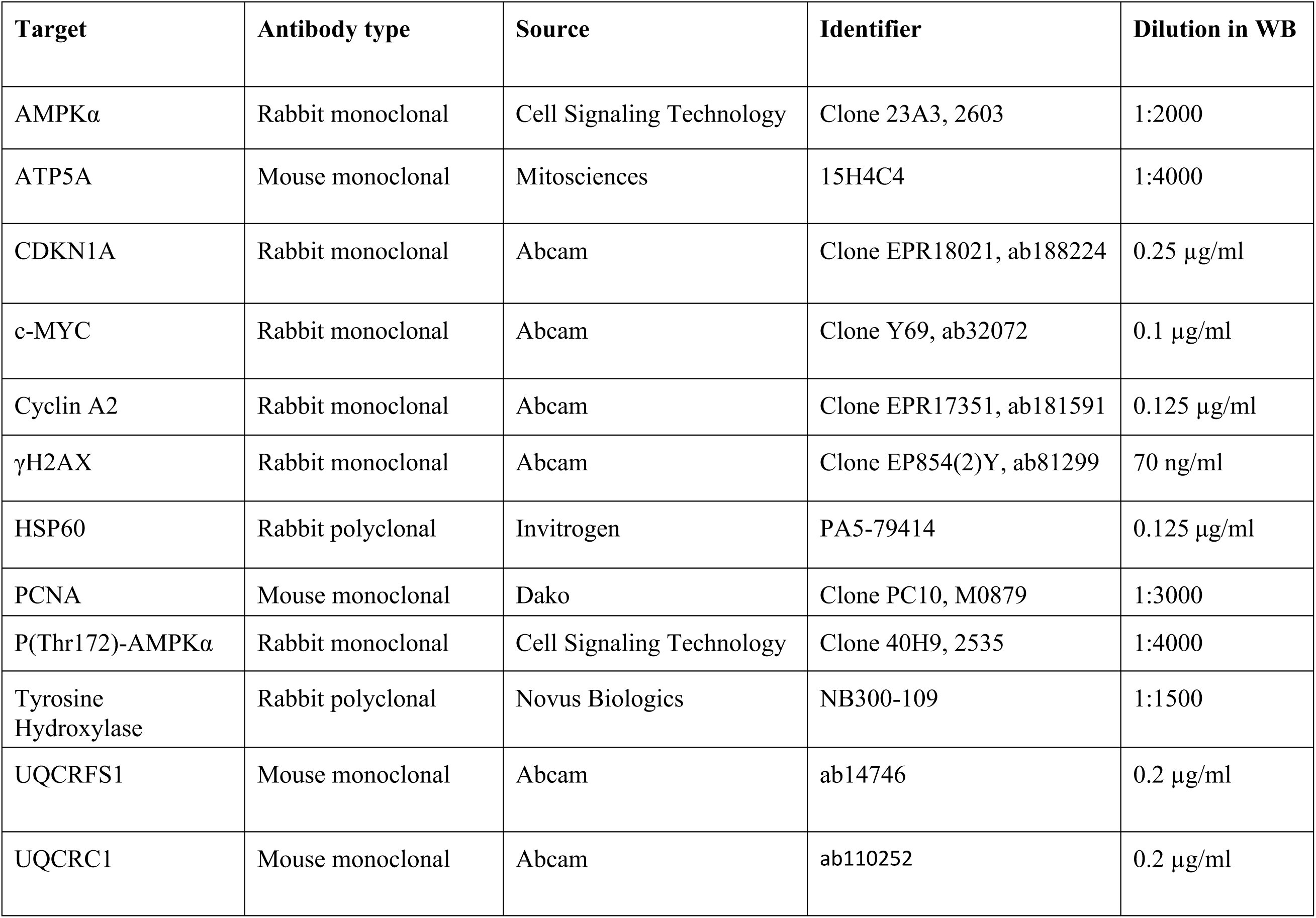
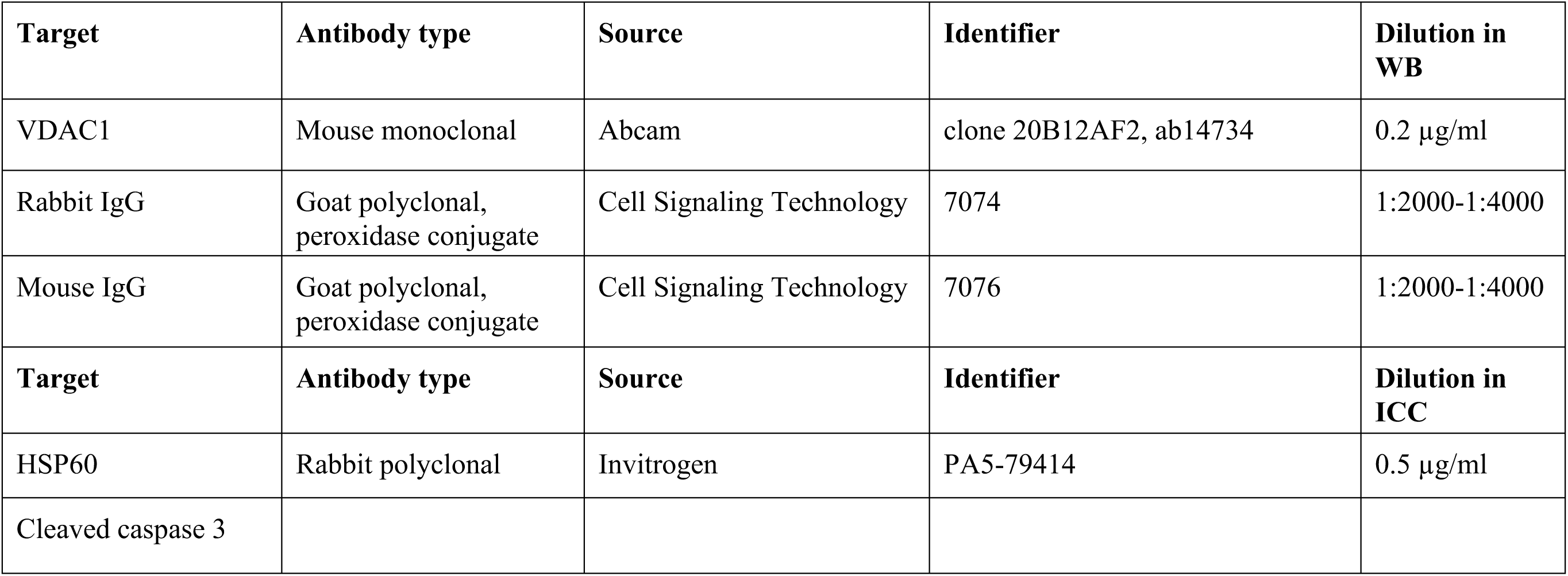
Antibodies.

